# Microscopic Quantification of Oxygen Consumption across Cortical Layers

**DOI:** 10.1101/2021.10.13.464176

**Authors:** Philipp Mächler, Natalie Fomin-Thunemann, Martin Thunemann, Marte Julie Sætra, Michèle Desjardins, Kıvılcım Kılıç, Ikbal Şencan, Baoqiang Li, Payam Saisan, Qun Cheng, Kimberly L. Weldy, David A. Boas, Richard B. Buxton, Gaute T. Einevoll, Anders M. Dale, Sava Sakadžić, Anna Devor

## Abstract

The cerebral cortex is organized in cortical layers that differ in their cellular density, composition, and wiring. Cortical laminar architecture is also readily revealed by staining for cytochrome oxidase – the last enzyme in the respiratory electron transport chain located in the inner mitochondrial membrane. It has been hypothesized that a high-density band of cytochrome oxidase in cortical layer IV reflects higher oxygen consumption under baseline (unstimulated) conditions. Here, we tested the above hypothesis using direct measurements of the partial pressure of O_2_ (pO_2_) in cortical tissue by means of 2-photon phosphorescence lifetime microscopy (2PLM). We revisited our previously developed method for extraction of the cerebral metabolic rate of O_2_ (CMRO_2_) based on 2-photon pO_2_ measurements around diving arterioles and applied this method to estimate baseline CMRO_2_ in awake mice across cortical layers. To our surprise, our results revealed *a decrease in baseline CMRO_2_ from layer I to layer IV.* This decrease of CMRO_2_ with cortical depth was paralleled by *an increase in tissue oxygenation.* Higher baseline oxygenation and cytochrome density in layer IV may serve as an O_2_ reserve during surges of neuronal activity or certain metabolically active brain states rather than baseline energy needs. Our study provides the first quantification of microscopically resolved CMRO_2_ across cortical layers as a step towards informed interpretation and modeling of cortical-layer-specific Blood Oxygen Level Dependent (BOLD) functional Magnetic Resonance Imaging (fMRI) signals.

## Introduction

Dramatic improvements in functional Magnetic Resonance Imaging (fMRI) technology in recent years, including significant advances in hardware and image reconstruction (Polimeni & Wald 2018, Ugurbil 2018, Wald 2012, Yu et al 2014), have enabled resolution of cortical layers bringing us one step closer to the spatial scale of local neuronal circuits (Dumoulin et al 2018, Goense et al 2016, Polimeni et al 2010, Turner 2016). Indeed, neuronal circuits in cerebral cortex are organized in layers that differ in their cellular density, composition and wiring (Helmstaedter et al 2007, Woolsey & Van der Loos 1970), as well as the density of mitochondrial cytochrome oxidase, a marker of O_2_ metabolism (Wong-Riley 1989). In primary cortices, the highest density of cytochrome oxidase is found in layer IV, and the lowest – in layer I (Land & Simons 1985, Weber et al 2008). Therefore, it is commonly believed that layer IV has higher cerebral metabolic rate of O_2_ (CMRO_2_) compared to other layers. These differences have been specifically hypothesized to reflect laminar variation in metabolic costs under the baseline (unstimulated) conditions (Weber et al 2008). Experimentally addressing this hypothesis is important for physiological interpretation of the Blood Oxygenation Level Dependent (BOLD) contrast used in cortical layer-resolved fMRI studies (Dumoulin et al 2018, Goense et al 2012, Hirano et al 2011, Siero et al 2015, Yu et al 2014).

Experimental measurement of layer-specific (i.e. laminar) CMRO_2_ has been challenging, because in common practice it requires information about both blood oxygenation and blood flow (Devor et al 2012a, Devor et al 2012b, Dunn et al 2005, Royl et al 2008). Previously, we introduced a method for extraction of CMRO_2_ (Sakadzic et al 2010) based on a single imaging modality: 2-photon phosphorescence lifetime microscopy (2PLM) (Finikova et al 2008, Lecoq et al 2011, Sakadzic et al 2010) providing measurements of the partial pressure of O_2_ (pO_2_). This method relies on the Krogh-Erlang model of O_2_ diffusion from a cylinder (Krogh 1919), which assumes that a volume of tissue within a certain radius around a blood vessel gets all its O_2_ from that blood vessel. Previously, we applied our method to estimate CMRO_2_ in the rat cerebral cortex, where capillaries were absent within an ~100-μm radius around penetrating arterioles satisfying the model assumption. The laminar profile of CMRO_2_, however, has not been addressed due to limited penetration of our imaging technology at that time.

In the present study, we increased our penetration depth by (i) utilizing a new pO_2_ probe Oxyphor 2P with red-shifted absorption and emission spectra (Esipova et al 2019), (ii) switching from rats to mice since mice have a thinner cortex and smaller diameter of surface blood vessels attenuating light, and (iii) optimizing our procedure of probe delivery to avoid spilling of the probe on the cortical surface exacerbating out-of-focus excitation. With these improvements, we performed pO_2_ measurements in fully awake mice in cortical layers I-IV.

In the mouse cortex, periarteriolar spaces void of capillaries are narrower than those in the rat (Kasischke et al 2011, Uhlirova et al 2016b). This challenges the basic Krogh-Erlang assumption of the center arteriole serving as the sole O_2_ source. Therefore, we revisited the model to account for the contribution of the capillary bed and then applied the revised model to quantify the baseline laminar CMRO_2_ profile. Our results show that, contrary to the common notion (Weber et al 2008), baseline CMRO_2_ in layer IV does not exceed that in the upper cortical layers. We speculate that higher cytochrome oxidase density in layer IV may reflect higher O_2_ metabolism during transient surges of neuronal activity or certain metabolically active brain states rather than baseline energy needs.

## Results

### Depth-resolved measurements of tissue pO_2_ in awake mice with chronic optical windows

We used 2PLM (**Fig. 1A**) (Finikova et al 2008, Lecoq et al 2011, Sakadzic et al 2010) in combination with a new pO_2_ probe Oxyphor 2P (Esipova et al 2019) to image baseline interstitial (tissue) pO_2_ in the primary cerebral cortex (SI) around diving arterioles at different depths: from the cortical surface down to ~500 μm deep. All measurements were performed in awake mice with chronically implanted optical windows (**Fig. 1B-C** and **Suppl Fig. 1**). Oxyphor 2P was pressure-microinjected into the cortical tissue ~400 μm below the surface and was allowed to diffuse resulting in sufficient signal intensity in cortical layers I-IV within ~40 min after the injection. Sulforhodamine 101 (SR101) was co-injected with Oxyphor 2P to label astrocytes as a means for real-time monitoring of tissue integrity. In addition, fluorescein isothiocyanate (FITC)-labeled dextran was injected intravenously to visualize the vasculature. We imaged pO_2_ using either square or radial grids of 400 points in the XY plane covering an area around a diving arteriole up to ~200 μm in the radial distance. Overall, pO_2_ measurements were acquired at 24 planes along 6 diving arterioles in 4 subjects.

**Figure 1.**
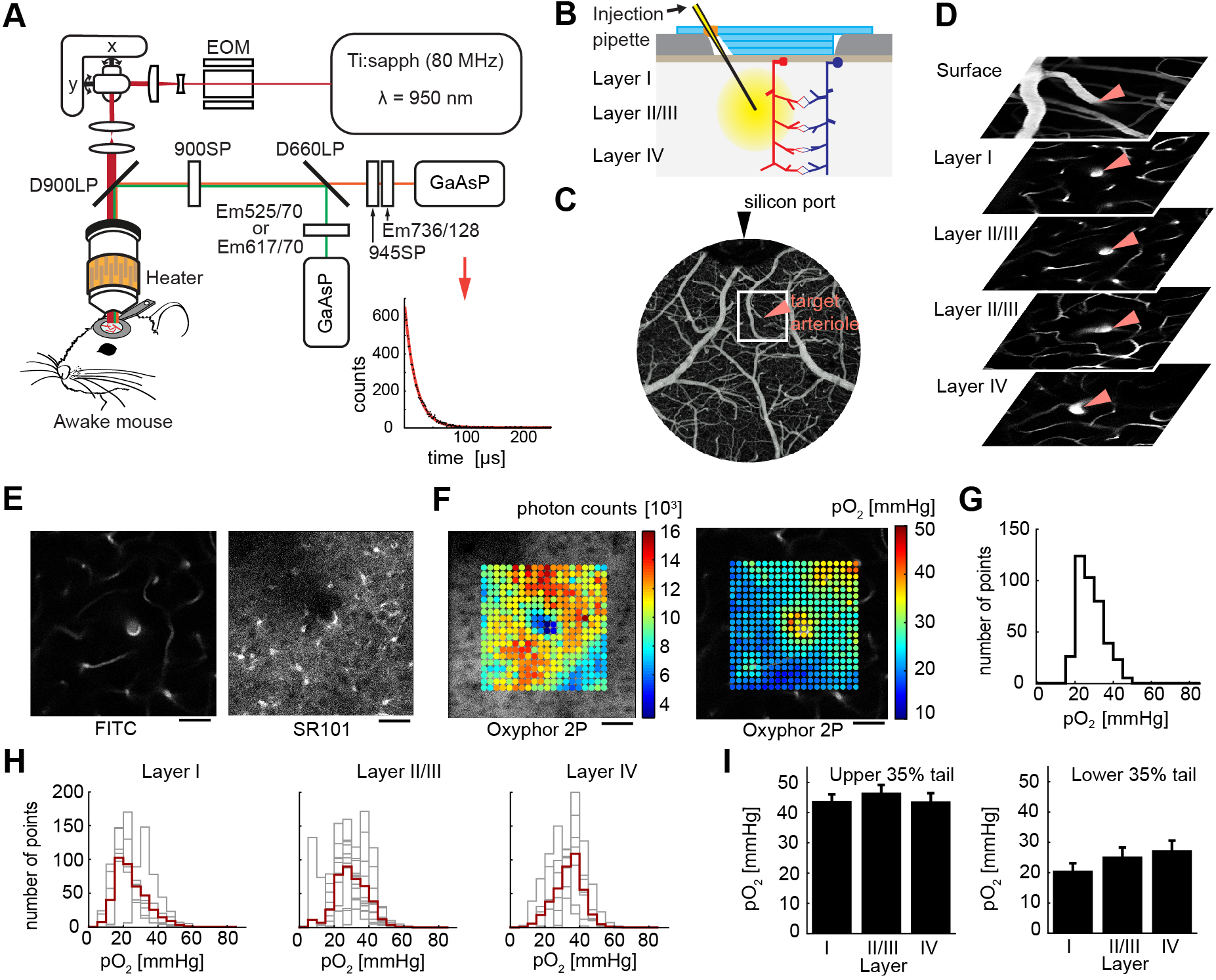
Measurements of tissue pO_2_ across cortical layers of awake mice using 2-photon phosphorescence lifetime microscopy (2PLM) **A.** Imaging setup for 2PLM in awake, head-restrained mice. Ti:sapph (80 MHz) – femtosecond pulsed laser tuned to 950 nm; EOM – electro-optic modulator; D900LP and D660LP-long pass dichroic mirrors with a cutoff at 900 and 660 nm, respectively; GaAsP – photomultiplier tubes; 900SP and 945SP – short pass optical filters with a cutoff at 900 and 945 nm, respectively; Em525/70, Em617/70, Em736/128 – bandpass emission filters. The inset in the lower right corner illustrates a phosphorescence decay; data (black) and fit (red) are overlaid. **B.** Schematics of the chronic cranial window with a silicon port for intracortical injection of Oxyphor 2P. **C.** An image of surface vasculature calculated as a maximum intensity projection (MIP) of a 2-photon image stack 0-300 μm in depth using a 5x objective. Individual images were acquired every 10 μm. Fluorescence is due to intravascular FITC. **D.** An example set of images tracking a diving arteriole throughout the top 500 μm of cortex (red arrows). Fluorescence is due to intravascular FITC. **E.** A measurement plane 200 μm deep including intravascular FITC (left) and SR101-labeled astrocytes (right) for the same arteriole as in (D). Scale bar = 50μm **G.** Histogram of pO_2_ values corresponding to (F). **F.** A square measurement grid of 20×20 points obtained from the imaging plane shown in (E). Left: photon counts are overlaid on the image of phosphorescence. Right: calculated pO_2_ values superimposed on a vascular FITC image. Scale bar = 50μm **H.** pO_2_ histograms across cortical layers. Each panel shows overlaid histograms from each measurement plane (gray) and superimposed average (red). **I.** Quantification of the top and bottom 35% of the pO_2_ distributions from (H). Error bars show standard error calculated using a mixed effect model implemented in *R* (p>0.1 for the top 35% and p<0.05 for the bottom 35%).

We traced individual diving arterioles from the cortical surface and acquired grids of pO_2_ points with an arteriole at the center (**Fig. 1D** and **Suppl Fig. 2**). **Figure 1E** shows an example imaging plane crossing a diving arteriole. For each plane, a corresponding “reference” vascular image of intravascular FITC fluorescence was acquired immediately before and immediately after the pO_2_ measurements for coregistration of the measurement points in the coordinate system of the vascular network. The measurements are color-coded according to the pO_2_ level (in mmHg) and superimposed on the reference vascular image (**Fig. 1F**).

We grouped pO_2_ measurements according to the following depth categories: layer I (50-100 μm), layer II/III (150-300 μm) and layer IV (320-500 μm). **Figure 1G** shows distributions of tissue pO_2_ values for the specific example shown in **Figure 1E-F**, and **Figure 1H** shows similar histograms for the overall dataset pooled across subjects. The high tail of the overall pO_2_ distribution (the upper 35%), reflecting oxygenation in tissue in the immediate vicinity of arterioles, did not change significantly as a function of depth. This lack of dependence of the peri-arteriolar pO_2_ on depth is in agreement with recent depth-resolved intravascular pO_2_ studies in awake mice by us and others showing only a small decrease in the intravascular pO_2_ along diving arteriolar trunks (Li et al 2019, Lyons et al 2016, Sencan et al 2020). In contrast, there was a significant increase in the low tail (the bottom 35%) of the pO_2_ distribution with depth (**Fig. 1I,** p<0.05) presumably indicating an increase in oxygenation of tissue in between capillaries in layer IV compared to layer I. This depth-dependent increase questions the common believe that layer IV has higher baseline CMRO_2_ compared to other layers. In addition, the median of the pO_2_ distribution shifted to the higher pO_2_ values below 200 μm (**Fig. 1H**). The median increase can be explained, at least in part, by branching: at certain depths, diving arterioles branched leading to the presence of highly oxygenated vessels (arterioles and capillaries) in the vicinity of diving arteriolar trunks.

### DACITI model for CMRO_2_ extraction

To quantify the laminar profile of CMRO_2_ in mouse cerebral cortex, we revised our model for estimation of CMRO_2_ based on peri-arteriolar pO_2_ measurements. The vascular architecture in the cerebral cortex features the absence of capillaries around diving arterioles (Kasischke et al 2011, Sakadzic et al 2014, Sakadzic et al 2016). Previously, we and others have argued that this organization agrees with the Krogh-Erlang model of O_2_ diffusion from a cylinder (Krogh 1919), where a diving arteriole can be modeled as a single O_2_ source for the tissue in the immediate vicinity of that arteriole where no capillaries are present. Indeed, radial pO_2_ gradients around cortical diving arterioles in the rat SI approached zero at the tissue radii corresponding to the first appearance of capillaries, and application of this model to estimate CMRO_2_ yielded physiologically plausible results (Sakadzic et al 2016). In the mouse, however, capillary-free peri-arteriolar spaces are smaller compared to those in the rat, and O_2_ delivery by the capillary bed cannot be ignored (Moeini et al 2019). Therefore, we revised the model to explicitly include both arteriolar and capillary delivery as we describe below.

In the Krogh-Erlang model, a blood vessel – approximated as an infinitely long cylinder – has a feeding tissue territory with a radius *r* = *R_t_* Outside of this cylinder, we enter a feeding territory of other vessels. Therefore, according to the Krogh-Erlang model, tissue pO_2_ will monotonically decrease with *r* while moving away from the vessel until we reach *R_t_* The minimum occurs at *r* = *R_t_* where the first spatial derivative along the radial direction is zero (*dpO_2_/dr* = 0). Outside of this cylinder, the predicted pO_2_ for *r* > *R_t_*. has no physiological meaning as the model is defined for *r* ≤ *R_t_*. However, Krogh-Erlang formula for *r* > *R_t_* predicts rapid pO_2_ increase, which is not compatible with the experimental observations of small tissue pO_2_ changes in the capillary bed and it complicates the fitting procedure (Sakadzic et al 2016). Importantly, in the cerebral cortex, not only arterioles but also capillaries contribute to tissue oxygenation, and this contribution is larger in awake animals (Li et al 2019, Sencan et al 2020) compared to those under anesthesia (Sakadzic et al 2014). However, radial pO_2_ changes around capillaries are still small (Li et al 2019, Lu et al 2020) and our current pO_2_ measurements are not optimal for accurately mapping them. Therefore, to account for the pO_2_ distribution in the capillary bed (*r* > *R_t_*), we revised the model assumptions as follows: (i) similar to the Krogh-Erlang model, we assumed that within a certain circular distance around a diving arteriole (*r* < *R_t_*), the arteriole in the center is the dominant source of O_2_ such that 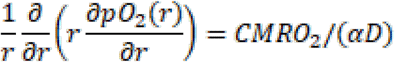, where *α* and *D* are O_2_ solubility and diffusivity, respectively; (ii) for (*r* > *R_t_*), we assume that the O_2_ supply from capillaries is uniformly distributed and equal to the O_2_ consumption, such that 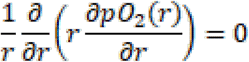. The new model, which we call ODACITI (**O**_2_ **D**iffusion from **A**rterioles and **C**apillaries **I**nto **TI**ssue), is schematically illustrated in **Figure 2A**. As for the Krogh-Erlang model, the central arteriole (red) with the radius *r* = *R_ves_* is feeding the peri-arterial tissue (**Fig. 2A**, pink) with the radius *r* = *R_t_*. Beyond that, O_2_ is supplied by the capillary bed so that the second spatial derivative is zero for *r* > *R_t_* (**Fig. 2A**, blue). With this set of assumptions, we derived a new specific solution to the forward Poisson problem in cylindrical coordinates (see **Methods**).

**Figure 2.**
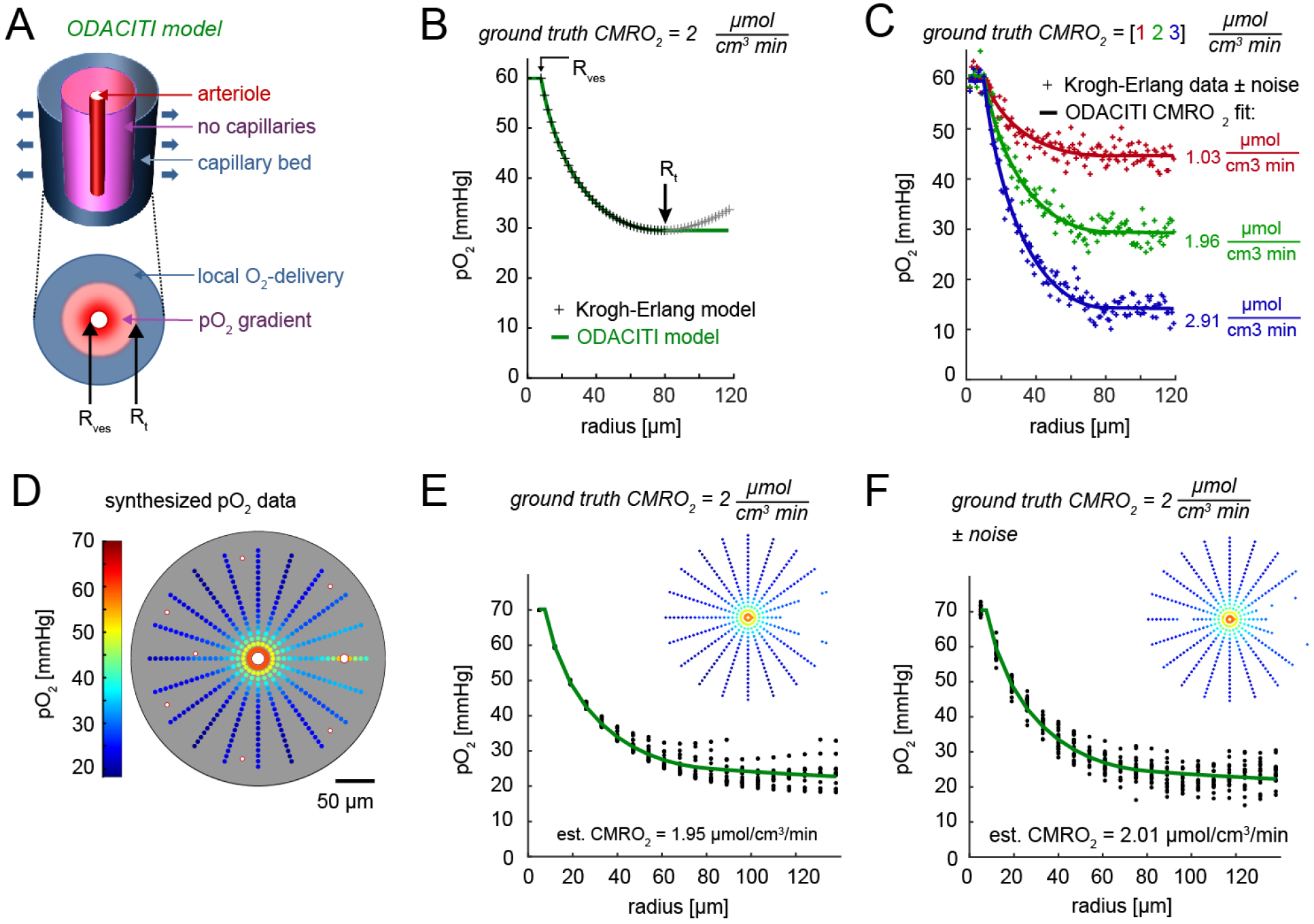
ODACITI model and validation with synthetic data. **A.** Schematic illustration of the model assumptions. The center arteriole is the only source of O_2_ for the pink peri-arteriolar region void of capillaries extending out to radius r=R_t_. For r>R_t_, delivery and consumption are balanced. **B.** Comparison of the functional form of the Krogh-Erlang (black) and ODACITI model (green).The Krogh-Erlang model but not ODACITI forces an increase in pO_2_ beyond R_t_. **C.** Application of ODACITI to synthetic data with added Gaussian noise (σ=2). In these data, pO_2_ = P_ves_ for r< R_ves_ and pO_2_ = pO_2_(R_t_) for r>R_t_. For R_ves_<r<R_t_, we solved for pO_2_ using the Krogh-Erlang equation. Three cases with CMRO_2_ of 1, 2 and 3 μmol cm-3 min-1 (color-coded) are superimposed. The ODACITI fit (lines) are overlaid on the data (points). **D.** Simulated pO_2_ data generated by solving the Poisson equation in 2D for a given geometry of vascular O_2_ sources, including one highly oxygenated vessel (on the right) and given CMRO_2_ =2 μmol cm-3 min-1. **E.** The pO_2_ gradient as a function of the distance from the center arteriole. The green line shows ODACITI fit. **F.** As in (E) after adding Gaussian noise (σ = 2 SD).

### Validation of the ODACITI model using synthetic data

To validate the ODACITI model, we used synthetic data where the ground truth (space-invariant) CMRO_2_ was preset and thus known. First, we compared the functional form of the radial pO_2_ gradient obtained with ODACITI to that obtained with the Krogh-Erlang equation. To that end, we generated synthetic data by analytically solving the Krogh-Erlang and ODACITI equation for a given CMRO_2_ and *R_t_*. As expected, the models agree for *r* < *R_t_* producing the same monotonic decrease in pO_2_ with increasing distance from the center arteriole. Beyond *R_t_*, the Krogh-Erlang formula produces an unrealistic increase in pO_2_, while ODACITI is able to take into account measured tissue pO_2_ in the capillary bed and it produces more realistic, nearly constant level of pO_2_ (**Fig. 2B**). Next, we tested ODACITI on synthetic data with added noise to mimic noise of experimental measurements. To that end, we (1) amended the Krogh-Erlang solution by imposing a constant pO_2_ equal to pO_2_(R_t_) for *r* > *R_t_* to represent the capillary bed, and (2) added Gaussian noise (σ=2). With these synthetic data, we were able to sufficiently recover the true CMRO_2_ with ODACITI (**Fig. 2C**). In another test, we performed fitting while selecting a constant *R_t_* between 60 – 80 μm. This resulted in an 18% error in the estimated CMRO_2_ (**Suppl. Fig. 3**). In comparison, the same variation in resulted in a 45% error while using a fitting procedure developed in Sakadzic et al. 2014 to fit for CMRO_2_ based on the Krogh-Erlang model (Sakadzic et al 2014). Thus, ODACITI is less sensitive to the error in *R_t_*. compared to the Krogh-Erlang model.

ODACITI assumes that the first spatial derivative *dpO*_2_(*r*)/*dr* monotonically decreases in the proximity of the arteriole and remains near zero in the capillary bed. In reality, however, this derivative varies in different directions from the center arteriole due to asymmetry of the vascular organization. Moreover, for some radial directions, pO_2_ may remain high due to the presence of another highly oxygenated vessel (a small arteriolar branch or highly oxygenated capillary (Li et al 2019, Sakadzic et al 2014). To model this situation, we placed additional O_2_ sources representing highly oxygenated blood vessels around the center arteriole (**Fig. 2D**). When multiple nearby vessels serve as O_2_ sources, no analytical solution for the tissue pO_2_ map is available. Therefore, we solved the Poisson equation, which relates CMRO_2_ and tissue pO_2_, by means of finite-element numerical modeling implemented in the software package FEniCS (Logg et al 2012) (see **Methods**). Previously, we verified this implementation (for simple vascular geometries) by comparing the result to that of the Krogh-Erlang equation (Saetra et al 2020). **Fig. 2D** illustrates an example FEniCS output for a center 15-μm-diameter “arteriole” surrounded by seven 5-μm-diameter “capillaries” located 80-130 μm away from the diving arteriole (**Fig. 2D**), which is a typical size of the region around diving arterioles void of capillaries in mouse cerebral cortex. The intravascular pO_2_ was set to 70 mmHg and 35 mmHg for the arteriole and capillaries, respectively. In addition, we placed another “arteriole” with intravascular pO_2_ was set to 60 mmHg 110 μm away from the center arteriole to simulate the occasional presence of highly oxygenated vessels within our measurement grid.

These simulated data were used to devise a procedure for segmenting a region of interest (ROI) consistent with the assumption that pO_2_ should decrease monotonically from the center arteriole until reaching a certain steady-state value within the capillary bed (see **Methods**). We then used the data included in the ROI to extract the pO_2_ profile as a function of the radial distance from the center arteriole. Finally, we applied the ODACITI model to these radial pO_2_ profiles to calculate CMRO_2_. We calculated CMRO_2_ in a noise-free case as well as with additive Gaussian noise (see **Methods**). In both cases, the model was able to recover the true CMRO_2_ (**Fig. 2E-F**).

### Estimation of CMRO_2_ across cortical layers

Following validation with synthetic data (**Fig. 2**), we used ODACITI to calculate CMRO_2_ across the cortical depth keeping the same depth categories as in **Figure 1**. For each imaging plane, we registered a pO_2_ measurement grid with the corresponding vascular reference image. As with synthetic data, we segmented an ROI for each measurement grid and collapsed the data within each ROI into a corresponding radial pO_2_ gradient (**Fig. 3A**, see **Methods**). These peri-arteriolar pO_2_ gradients are consistent with our previous measurements in the rat cortex under α-chloralose anesthesia (Devor et al 2011) and a more recent study in superficial cortex of awake mice (Lu et al 2020). Next, we performed fitting of the data to the ODACITI solution to quantify CMRO_2_ (Fig. 3B) (see **Methods**). **Figure 3C** shows laminar (depth-resolved) CMRO_2_ profiles along 6 diving arterioles in 4 subjects. We used a mixed effects model implemented in *R* to quantify CMRO_2_ across the layers. This analysis revealed a significant reduction of CMRO_2_ with depth (p=0.001, **Fig. 3D**). For this calculation, *R_t_* was fixed at 80 μm (see **Methods**); allowing *R_t_* to vary while fitting for CMRO_2_ did not reveal dependence of *R_t_* on cortical depth (not shown).

**Figure 3.**
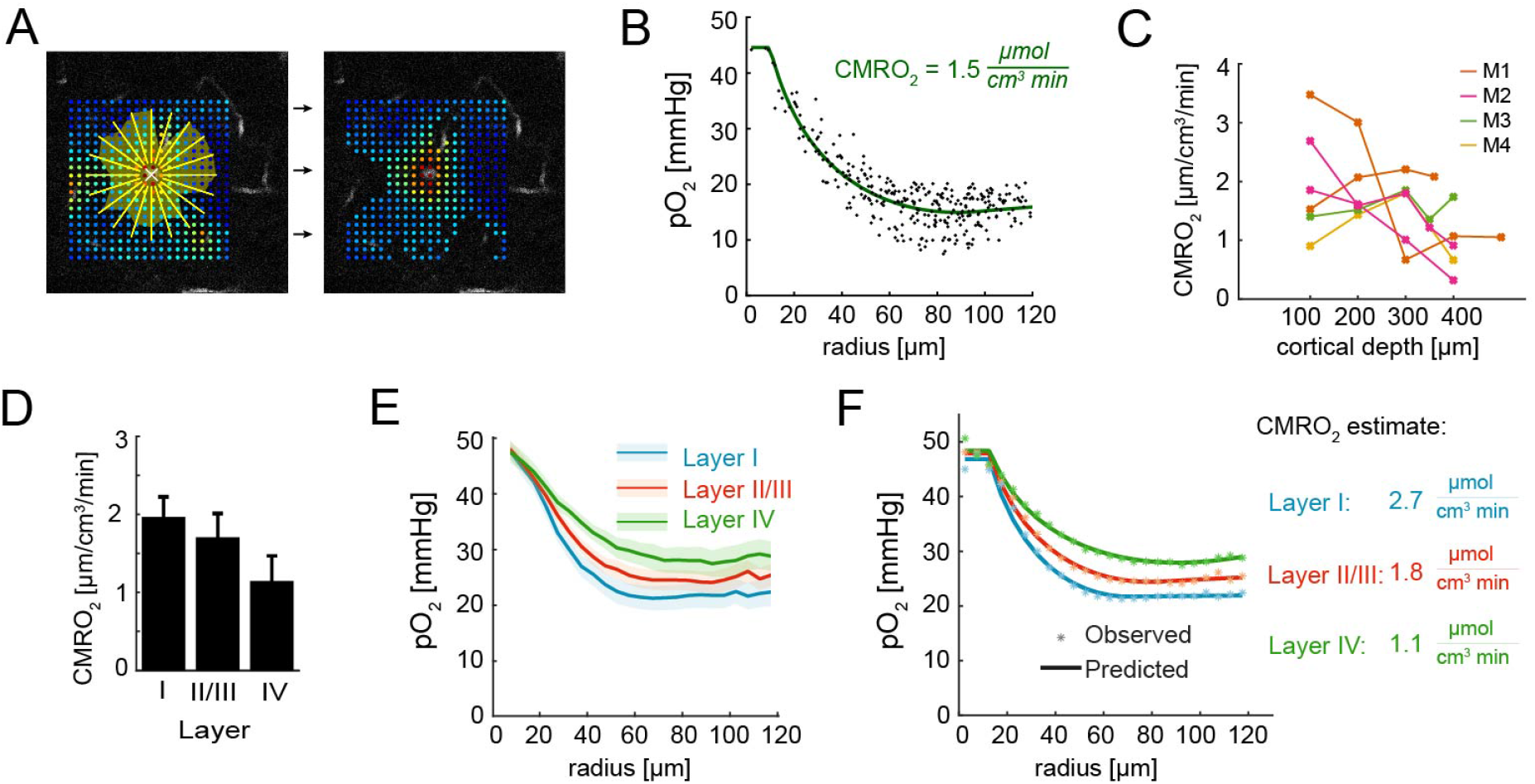
CMRO_2_ estimation across cortical layers. **A.** An imaging plane 100 μm below the surface; a grid of pO_2_ points is overlaid on the vascular image. pO_2_ values were interpolated along the radial yellow lines. The yellow shaded area extends along each direction until the first derivative becomes zero. Segmented ROI used for extraction of the radial pO_2_ profile is shown on the right. This ROI includes in addition values within the low tail of the pO_2_ distribution. **B.** The radial pO_2_ profile extracted from (A). ODACITI fit is overlaid on the data. The fitted CMRO_2_ value is 1.5 μmol cm^-3^ min^-1^. **C.** Estimated CMRO_2_ for the entire dataset of 24 planes along 6 diving arterioles in 4 subjects. Measurements along the same arteriole are connected in a line. Subjects are color-coded. **D.** Quantification of CMRO_2_ across layers using the data in (C). Error bars show standard error calculated using a linear mixed effect model implemented in *R.* **E.** Superimposed color-coded grand averaged radial pO_2_ profiles for layers I, II/III and IV (mean±SEM for 5-μm bins). For each layer, the profile was obtained by averaging all measurements within that layer across arterioles and subjects. **F.** CMRO_2_ extracted from averaged profiles in (E).

For each arteriole, we also acquired pO_2_ measurements at the cortical surface (**Suppl Fig. 4**). The surface measurements, however, were not used for estimation of CMRO_2_ because of violation of the axial symmetry assumption (i.e., absence of cerebral tissue above the imaging plane).

Graphically, CMRO_2_ scales with the steepness of the descent of peri-arteriolar radial pO_2_ profile (Kasischke et al 2011, Krogh 1919, Saetra et al 2020, Sakadzic et al 2016). In **Figure 3E**, we overlay grand averaged radial profiles for each layer. Each curve was obtained by averaging all data for that layer (**Suppl Fig.5**). As can be appreciated by visual inspection, the gradient corresponding to layer I has the steepest descent, while the descent of gradient in layer IV is more relaxed. This indicates that CMRO_2_ in layer IV is lower compared to the upper layers, which is consistent with the model-based estimation shown in **Figure 3D**.

Taken together, our results indicate that an increase in the low tail of the pO_2_ distribution with depth, which likely reflects an increase in oxygenation of tissue in between capillaries in layer IV compared to upper layers (**Fig. 1H-I**), can be explained at least in part by a depth-dependent decrease in CMRO_2_ from layer I to layer IV (**Fig. 3D-F**). This is in contrast to a well-established fact that capillary, cytochrome oxidase, and mitochondrial densities in the mouse SI all peak in layer IV (Blinder et al 2013, Santuy et al 2018) implying that neither of these densities can serve as a proxy for oxidative energy metabolism under baseline (unstimulated) conditions.

## Discussion

In this work, we have achieved depth-resolved tissue pO_2_ measurements in fully awake mice using 2PLM in combination with the O_2_-sensitive probe Oxyphor-2P. We devised the ODACITI model for estimation of CMRO_2_ from peri-arteriolar pO_2_ gradients accounting for both arteriolar and capillary O_2_ supply. We validated the model using synthetic data and then applied it to estimate the laminar CMRO_2_ profile during the baseline level of neuronal activity (i.e., in the absence of external stimulation). Our results demonstrate a significant reduction of CMRO_2_ with depth from layer I to layer IV. While these results are based on a model and its specific assumptions, they strongly suggest that, at the very least, the baseline CMRO_2_ in layer IV does not exceed that in the upper cortical layers.

In cerebral cortex, neuronal cell type distribution as well as cellular, synaptic and microvascular densities vary between cortical layers (Blinder et al 2013, Feldmeyer et al 2013, Harris & Shepherd 2015, Helmstaedter et al 2007, Hyder et al 2013, Li et al 2019, Markram et al 2004, Weber et al 2008, Wu et al 2016). Cortical laminar architecture is also readily revealed by counting mitochondria (Santuy et al 2018) or staining for cytochrome oxidase – the last enzyme in the respiratory electron transport chain located in the mitochondrial membrane (Weber et al 2008, Wong-Riley 1989). In SI, cytochrome oxidase distribution features a high-density band in cortical layer IV, which has been hypothesized to reflect higher CMRO_2_ compared to other layers. Further, cytochrome oxidase was found to have a strong correlation with microvascular but not synaptic density – a finding that was interpreted to reflect high layer IV CMRO_2_ in the “idling state” (Weber et al 2008), where about half of the energy expenditure reflects other processes than synaptic and spiking activity (Attwell & Laughlin 2001, Engl & Attwell 2015). The present measurements and calculations do not support the idea of high baseline CMRO_2_ in layer IV.

If so, what is the purpose of high cytochrome oxidase density in layer IV? In our recent study, we found that intravascular pO_2_ changes (ΔpO_2_) in response to a sensory stimulus across layers were conserved (Sencan et al 2020) indicating a conserved ratio between the demand and supply. Taken together with existing data on larger stimulus-induced increases in CBF (ΔCBF) in layer IV compared to other layers (Srinivasan & Radhakrishnan 2014), this finding implies that the laminar ΔCMRO_2_ profile should also peak in layer IV in order for ΔpO_2_ to remain invariant. Thus, cytochrome oxidase may reflect layer-specific differences in peak energetic demands during transient neuronal signaling events. These events may include not only task-induced computation performed by local neuronal circuits (e.g., Barrel cortex response to a whisker touch (Feldmeyer et al 2013)) but also other dynamic neuronal processes occurring on larger spatiotemporal scales such as neuromodulation and sleep (Madsen et al 1991).

In this study, we observed a shift towards higher oxygenation of tissue within the capillary bed – quantified as the mean of the low tail of pO_2_ distribution – with increasing depth. Because our pO_2_ grids were always placed around diving arterioles, the low tail of the pO_2_ distribution may not reflect oxygenation of the capillary bed remote from arteriolar O_2_ sources. This observation, however, is potentially in line with our recent intravascular pO_2_ study in awake mice, where we showed that mean capillary pO_2_ in layer IV was ~15% higher than that in the upper layers (Li et al 2019). It is also consistent with a recent study where tissue pO_2_ measurements were performed at different depths while a Clark-type polarographic electrode was advanced throughout cortical layers (Zhang et al 2019). We speculate that higher baseline oxygenation in layer IV may serve as an O_2_ reserve for transient increases in neuronal activity. Beyond this reserve, upregulation in aerobic glycolysis may play a role alongside oxidative phosphorylation to rapidly supply ATP (Dienel 2019, Raichle & Mintun 2006, Yellen 2018).

In principle, higher capillary density in layer IV (Blinder et al 2013) may effectively shrink the peri-arteriolar cylinder where all O_2_ is provided by the diving arteriole. This hypothetical scenario, however, would not affect the pO_2_ gradient around the arteriole (which reflects CMRO_2_). Rather, it would decrease the radius of this cylinder, *R_t_*. Although in our calculations of CMRO_2_ *R_t_*. was a fixed parameter, we show that sensitivity of ODACITI to varying *R_t_* was relatively low. In addition, allowing *R_t_* to vary while fitting for CMRO_2_ did not reveal depth-dependence.

The current study is part of our ongoing effort to improve 2PLM (Devor et al 2011, Devor et al 2013, Esipova et al 2019, Li et al 2019, Sakadzic et al 2010, Sakadzic et al 2016, Sencan et al 2020) and microscopic estimation of CMRO_2_ (Saetra et al 2020, Sakadzic et al 2016, Uhlirova et al 2016a). In principle, if we knew space-resolved tissue pO_2_ as well as intravascular pO_2_ for each blood vessel within that tissue, we could solve for CMRO_2_ (that may vary in space and time) by finding an inverse solution for the Poisson diffusion equation (Saetra et al 2020). In practice, however, the signal to noise ratio (SNR) of our measurements may not be sufficient to accurately resolve pO_2_ gradients around capillaries. In addition, our current spatial resolution limits simultaneous measurements of capillary and tissue pO_2_, and sampling pO_2_ in 3D also remains a challenge. To mitigate these limitations, similar to the Krogh-Erlang model, the ODACITI model relies on radial pO_2_ gradients around diving arterioles assuming cylindrical symmetry. In reality, highly oxygenated arteriolar branches and/or low branching order capillaries can often be present on one side of a diving arteriole requiring ROI segmentation. Applied to synthetic data, our data analysis stream – including the segmentation algorithm followed by ODACITI estimation – was able to recover the “ground truth” CMRO_2_ validating the method. In the future, further improvements in pO_2_ probes and 2PLM technology may enable estimation of CMRO_2_ in 3D using numeric methods without the need to segment the data (Saetra et al 2020).

Our measurements were limited to cortical layers I-IV. In 2-photon microscopy, depth penetration is fundamentally limited by out-of-focus excitation (Theer & Denk 2006). Because light is scattered and absorbed by tissue, the laser power is ought to increase with depth to overcome these effects and deliver a sufficient number of photons to the focal volume. In our experiment, an increase in the number of scattered photons would increase the probability of out-of-focus excitation of Oxyphor 2P generating background phosphorescence. The lifetime of this background phosphorescence would report pO_2_ at locations outside the focal volume contributing to the experimental error and smoothing out peri-arteriolar pO_2_ gradients. In the future, a combination of 2PLM with light sculpting techniques (Rupprecht et al 2015) can be used to experimentally measure the amount of generated out-of-focus signal. 2PLM can also be implemented using two laser beams of different color that can be displaced in space minimizing out-of-focus excitation (Cheng et al 2020, Sadegh et al 2019).

This study was performed using fully awake mice with no anesthesia or sedation. While our mice were well trained to accept the head restraint and habituated to the imaging environment, we cannot rule out upregulation of adrenergic activity associated with arousal and/or stress. Previous studies have shown that aerobic glycolysis and glycogenolysis are triggered by activation of β1 and β2 adrenergic receptors, respectively (Dienel 2019, Dienel & Cruz 2016, Zuend et al 2020). Both of these processes are layer-specific in cerebral cortex. Therefore, the relative contribution of oxidative and glycolytic pathways to overall energy metabolism across cortical layers can depend on the state of adrenergic neuromodulation (Machler et al 2021). In the future, a combination of 2PLM with imaging of novel genetically encoded probes for norepinephrine and other neuromodulators (Feng et al 2019, Sabatini & Tian 2020) should allow quantification of CMRO_2_ in context of different brain states.

Knowing layer-specific CMRO_2_ in cerebral cortex is important for better understanding of the normal brain physiology as well as pathophysiology in diseases that affect cerebral microcirculation (Berthiaume et al., 2018; Iadecola, 2016, 2017; Pantoni, 2010; Zlokovic, 2011). It also is important for interpretation and modeling of the BOLD fMRI signals that are affected by both CBF and CMRO_2_ (Buxton 2010, Gagnon et al 2015, Uhlirova et al 2016a). Of particular relevance to the current results, the BOLD response is sensitive not only to the balance of the ΔCBF and ΔCMRO_2_ but also to their baseline state that can vary with age, disease, or even after consuming a cup of coffee (Griffeth et al 2011, Perthen et al 2008) altering the hemodynamic response for the same neuronal reality. Our study provides the first quantification of baseline CMRO_2_ across cortical layers as a step towards informed interpretation and modeling of high resolution BOLD signals.

## Methods

### O_2_ probe

The synthesis and calibration of the O_2_ probe Oxyphor 2P were performed as previously described (Esipova et al 2019). The phosphorescence lifetime of Oxyphor 2P is inversely proportional to the O_2_ concentration. Compared to its predecessor PtP-C343 (Finikova et al 2008), the optical spectrum of Oxyphor 2P is shifted to the longer wavelengths with the 2-photon absorption and phosphorescence peaks at 960 nm and 760 nm, respectively, enabling deeper imaging. Other advantages include large 2-photon absorption cross section (~600 GM near 960 nm) and higher phosphorescence quantum yield (0.23 in the absence of O_2_). In addition, the phosphorescence decay of Oxyphor 2P can be more closely approximated by a single-exponential function. Calibration for conversion of the phosphorescence lifetime to pO_2_ was obtained in independent experiments, where pO_2_ was detected by a Clark-type electrode as previously described (Esipova et al 2019).

### Two-photon imaging

Images were obtained using an Ultima 2-photon laser scanning microscopy system (**Fig. 1A**) (Bruker Fluorescence Microscopy). Two-photon excitation was provided by a Chameleon Ultra femtosecond Ti:Sapphire laser (Coherent) tuned to 950 nm.

For 2-photon phosphorescence lifetime microscopy (2PLM), laser power and excitation gate duration were controlled by two electro-optic modulators (EOMs) (350-160BK, Conoptics) in series with an effective combined extinction ratio > 50,000. This was done to mitigate bleed-through of the laser light and reduce the baseline photon count (i.e., during gate-OFF periods), which was critical for accurately fitting phosphorescence decays.

We used a combination of Zeiss 5x objective (Plan-NEOFLUAR, NA=0.16) for a coarse imaging and Olympus 20x (UMPlanFI, NA=0.5) objective for fine navigation under the glass window and for 2PLM. An objective heater (TC-HLS-05, Bioscience Tools) was used to maintain the temperature of the water between the objective lens and cranial window at 36.6 °C to avoid the objective acting as a heat sink and to comply with the Oxyphor 2P calibration temperature (Li et al 2019, Podgorski & Ranganathan 2016, Roche et al 2019).

The emission of Oxyphor 2P was reflected with a low-pass dichroic mirror (900-nm cutoff wavelength; custom-made by Chroma Technologies), subsequently filtered with a 795/150-nm emission filter (Semrock) and detected using a photon counting photomultiplier tube (PMT, Hamamatsu, H7422P-50). A second PMT, operated in analog mode (H7422-40, Hamamatsu) with 525/50 nm or 617/73 nm emission filters (Semrock) was used to detect fluorescein isothiocyanate (FITC)-labeled dextran and astrocytic marker sulforhodamine 101 (SR101), respectively (see *Delivery of optical probes* below). A 950-nm short-pass filter (Semrock) was positioned in front of the PMTs to further reject laser illumination.

At each imaging plane up to 400 μm below the cortical surface, a ~300 x 300 μm field of view (FOV) was selected for pO_2_ measurements. At each plane, pO_2_ measurements were performed serially arranged in a square or radial grid of 400 points.

The phosphorescence was excited using a 13-μs-long excitation gate, and the emission decay was acquired during 287 μs. The photon counts were binned into 2-μs-long bins. Typically, 50 decays were accumulated at each point with the total acquisition time of 15 ms per point. With 400 points per grid, one grid was acquired within 6 s. All points were revisited 20 times yielding a total of 1000 excitation cycles per point (50 cycles x 20 repetitions) acquired within 120 s. Dividing acquisition of 1000 cycles per point in blocks of 20 repetitions increased tolerance against motion (see *Motion correction* below) at a price of a slightly reduced sampling rate due to settling time of the galvonometer mirrors while moving from point to point.

### Implantation of the cranial window and headpost

All animal procedures were performed in accordance with the guidelines established by the Institutional Animal Care and Use Committee (IACUC) at University of California, San Diego. We used 13 adult C57BL/6J mice of either sex (age: 4-7 months). Seven of them were rejected due to imperfect healing of the cranial window, and 6 were used for pO_2_ measurements. Mice had free access to food and water and were held in a 12 h light/12 h dark cycle.

The surgical procedure for implantation of a chronic optical window was performed as previously described (Desjardins et al 2019, Kilic et al 2020). Briefly, dexamethasone was injected ~2 h prior to surgery. Mice were anesthetized with ketamine/xylazine or isoflurane (2% in O_2_ initially, 1% in O_2_ for maintenance) during surgical procedures; their body temperature was maintained at 37 °C. A 3-mm cranial window with a silicone port was implanted over the left barrel cortex, and the headpost was mounted over the other (right) hemisphere. The glass implant contained a hole, filled with silicone (Kilic et al 2020, Roome & Kuhn 2014), allowing intracortical injection of the O_2_ probe (**Fig.1B** and **Suppl. Fig. 1**). The window and the headpost were fixed to the skull in a predetermined orientation such that, when the mouse head was immobilized in the mouse holder (“hammock”), the window plane would be horizontal. Dextrose saline (5% dextrose, 0.05 ml) was injected subcutaneously before discontinuing anesthesia. Post-operative analgesia was provided with buprenorphine (0.05 mg/kg subcutaneously) injected ~20 min before discontinuing anesthesia. A combination of sulfamethoxazole/trimethoprim (Sulfatrim) (0.53mg/mL sulfamethoxazole and 0.11 mg/mL trimethoprim), and ibuprofen (0.05 mg/ml) was provided in drinking water starting on the day of surgery and for 5 days after surgery. Generally, full recovery and return to normal behavior were observed within 48 h post-op.

### Delivery of optical probes

Oxyphor 2P and astrocytic marker SR101 were delivered by intracortical microinjection through the silicon port in the glass window (**Fig. 1B** and **Suppl. Fig. 1**). Oxyphor 2P was diluted to 340 μM in artificial cerebrospinal fluid (ACSF, containing: 142 mM NaCl, 5 mM KCl, 10 mM glucose, 10 mM HEPES (Na Salt), 3.1 mM CaCl_2_, 1.3 mM MgCl_2_, pH 7.4) and filtered through a 0.2-μm filter (Acrodisc 4602, PALL). SR101 (Sigma S359) was added for a final concentration of 0.2 mM. We used quartz-glass capillaries with filament (O.D. 1.0 mm, I.D. 0.6 mm; QF100-60-10; Sutter Instrument). Pipettes were pulled using a P-2000 Sutter Instrument Puller. For each pulled pipette, the tip was manually broken under the microscope to obtain an outer diameter of 15-25 μm. A pipette was filled with a mixture of Oxyphor 2P and SR-101 (thereafter referred to as the “dye”) and fixed in a micromanipulator at an angle of ~35°. The pipette was guided under visual control through the silicon port to its final location in tissue while viewing the SR101 fluorescence through the microscope eyepiece. First, to target the tissue surrounding a diving arteriole, the pipette was oriented above the window such that its projection onto the window coincided with the line connecting the diving point with the silicon port. The pipette was then retracted and lowered to touch the surface of the port. There, a drop of the dye solution was ejected to ensure that the pipette was not clogged. A holding positive pressure was maintained (using PV830 Picopump, World Precision Instruments) to avoid clogging of the pipette while advancing through the port. An experimenter could recognize the pipette emerging below the port by the dye streaming from the tip. At this point, the holding pressure was quickly set to zero in order to avoid spilling of the dye on the cortical surface. Next, the pipette was advanced for ~600 μm along its axis at 35°. Below ~100 μm, some holding pressure was applied to allow leakage of the dye that was used to visualize the pipette advancement. The pressure was manually adjusted to ensure visible spread of the dye without displacing cortical tissue. When the target artery was reached, the pipette was withdrawn by ~50 μm, and holding pressure was maintained for ~20 min to allow slow diffusion of the dye into the tissue. A successful injection was recognized by the appearance of SR101 in the perivascular space around the targeted arteriole but not around its neighbors. The contrast between the targeted and neighboring arterioles was lacking when the dye was injected too shallow or too vigorously, in which case the experiment was aborted. After the loading, the pressure was set to zero and the pipette was withdrawn. The mouse was placed back in its home cage to recover for ~40 minutes until the start of the imaging session. At that time, the dye usually diffused within a 300-600-μm radius around the targeted arteriole.

FITC-dextran (FD2000S, 2 MDa, Sigma Aldrich) was used to visualize cortical vasculature. Mice were briefly anesthetized with isoflurane, and 50 μl of the FITC solution (5% in normal saline) was injected retro-orbitally prior to intracortical delivery of Oxyphor 2P for each imaging session.

### Habituation to head fixation

Starting at least 7 days after the surgical procedure, mice were habituated in one session per day to accept increasingly longer periods of head restraint under the microscope objective (up to 1 h/day). During the head restraint, the mouse was placed on a hammock. A drop of diluted sweetened condensed milk was offered every 20-30 min during the restraint as a reward. Mice were free to readjust their body position and from time to time displayed natural grooming behavior.

### Estimation of pO_2_

All image processing was performed using custom-designed software in MATLAB (MathWorks). For each point, cumulative data from all excitation cycles available for that point were used for estimation of the phosphorescent decay. Starting 5 μs after the excitation gate to minimize the influence of the instrument response function, the decay was fitted to a single-exponential function:
*****

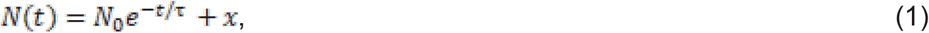

where *N*(*t*) is the number of photons at time *t*, *N*_0_ is the number of photons at *t*=0 (i.e., 5 μs after closing the excitation gate), *τ* is the phosphorescence lifetime and *x* is the offset due to non-zero photon count at baseline. The fitting routine was based on the nonlinear least-squares method using the MATLAB function *lsqnonlin.* The phosphorescence lifetime *τ* was then converted into absolute pO_2_ using an empirical biexponential form:

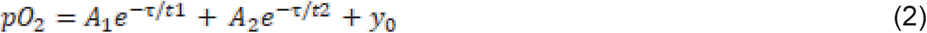

Where parameters *A*_1_, *t*_1_, *A*_2_, *t*_2_ and *y*_0_ were obtained during independent calibrations (Esipova et al 2019).

The locations of pO_2_ measurements were co-registered with FITC-labeled vasculature. Color-coded pO_2_ values are overlaid on vascular images (grayscale) in all figures displaying pO_2_ maps.

### Identification of data corrupted by motion

To exclude data with excessive motion, an accelerometer (ADXL335, Sainsmart, Analog Devices) was attached below the mouse hammock. The accelerometer readout was synchronized with 2-photon imaging and recorded using a dedicated data acquisition system (National Instruments). During periods with extensive body movement (e.g., grooming behavior), the accelerometer signal crossed a pre-defined threshold, above which data were rejected. In pilot experiments, in addition to the accelerometer, a webcam (Lifecam Studio, Microsoft; IR filter removed) with IR illumination (M940L3-IR (940 nm) LED, Thorlabs) was used for video recording of the mouse during imaging. The video and accelerometer reading were in general agreement with each other. Therefore, accelerometer data alone were used to calculate the rejection threshold. Because every point was revisited 20 times, typically at least 10 repetitions (or 500 excitation cycles) were unaffected by motion and were used to estimate τ. The MATLAB function *lsqnonlin* used to fit phosphorescent decays returned the residual error for each fit, that we plotted against the number of cycles (**Suppl. Fig. 6**). At around 500 cycles, the error stabilized at a low level. Therefore, we quantified points where at least 500 cycles were available.

### Estimation of CMRO_2_

The O_2_ transport to tissue is thought to be dominated by diffusion. In the general case, the relationship between pO_2_ denoted as 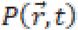 and the O_2_ consumption rate denoted *CMRO*_2_ (*r,t*) can be described by

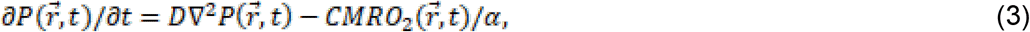

where ∇^2^ is the Laplace operator in three spatial dimensions, and *D* and *α* are diffusion coefficient and solubility of O_2_ in tissue, respectively. For all our calculations, we assumed α = 1.39 μM mmHg^-1^ and D = 4×10^-5^ cm^2^ s^-1^ (Goldman 2008). This equation is only applicable outside the blood vessels supplying O_2_ to the tissue. Oxygen supplied by a blood vessel is represented by a boundary condition of pO_2_ imposed at the vessel wall. When the system is in steady state, the term *∂P*(*r,t*)/*∂t* can be neglected. If we also assume that there is no local variation of pO_2_ in the vertical z-direction, that is, the direction along the cortical axis parallel to penetrating arterioles, **Eq. 3** simplifies to

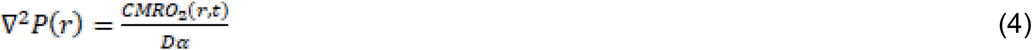

where *P*(*r*) represents pO_2_ measured at the radial location *r* and ∇^2^ refers to the twodimensional Laplace operator.

Here, we derive a specific solution to the forward problem of this partial differential equation using the following assumptions: First, as with the Krogh-Erlang model (Krogh 1919), we assume that tissue CMRO_2_ in the vicinity of a diving arteriole is space-invariant. Second, we assume that within a capillary-free region around the arteriole, all O_2_ is provided by that arteriole. That means that if we have an arteriole with the radius r=R_ves_ and a region around the arteriole void of capillaries within a radius r=R_t_, for the radial distance R_ves_<r<R_t_, the rate of O_2_ consumption is equal to CMRO_2_ (that we are trying to find). This is because no local O_2_ sources are present within this region except for the arteriole in the center. Third, we assume that for r>R_t_ the rate of O_2_ delivery by the capillary bed is uniformly distributed and equal to the rate of tissue O_2_ consumption. With these assumptions, we can derive an analytic solution to **Eq. 4** that we describe below.

We generally denote the partial differential equation in cylindrical coordinates as

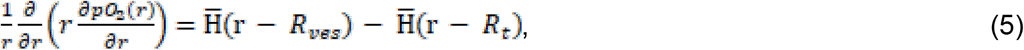

where *pO*_2_(*r*) is the pO_2_ as a function of the radial distance from the center of a diving arteriole, and *H*(*r*) is a Heaviside step function 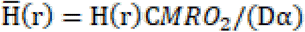. Therefore, Eq. 5 considers three spatial regions 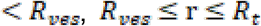, and r < *R_t_*, where only in the middle region (e.g., for *R_ves_* ≤ *r* ≤ *R_t_*) the right hand side of Eq. 5 is non zero. Additionally, we required that *pO*_2_(*r*) is a continuous function at r = *R_t_* (e.g.,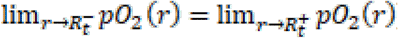) and at r = *R_t_* (e.g., 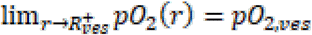). (e.g., 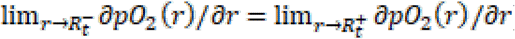). Importantly, to better account for the O_2_ delivery by the capillary bed that surrounds the arteriole, we assumed that *∂pO*_2_(*r*)/*∂r* = 0 for some r = *R*_0_, where *R_ves_* < *R*_0_ ≤ *R^t^* Finally, pO_2_ values that we measured at *r* = *R_ves_* did not show any diffusion gradients because they originated from extracellular dye in the tissue next to the vessel. Therefore, in our model we assumed that 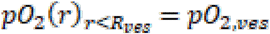. We considered the following solution of Eq. 5 which satisfies the above conditions:

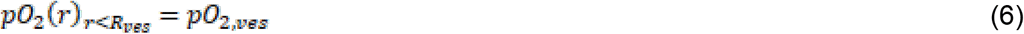

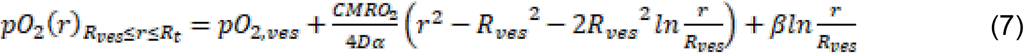

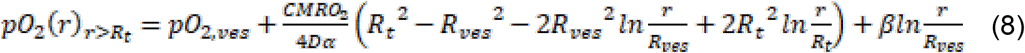

This model, which we nicknamed “ODACITI”, for **O**_2_ **D**iffusion from **A**rterioles and **C**apillaries **I**nto **TI**ssue, was used to fit for *CMRO*_2_,*pO*_2,*ves*_, and *β*, given the arteriolar radius *R_ves_* and the radius of capillary-free periarteriolar space *R_t_*. *R_ves_* was estimated from vascular FITC images as the full-width at half-maximum of the intensity profile drawn across the arteriole. was fixed to 80 μm based on observation of the capillary bed in mice. To simplify the fitting procedure, parameter *β* was used in Eqs. 6–8 to replace 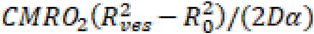.

Notably, ODACITI model behaves not necessarily the same at Rt as the original Krogh-Erlang model given by:

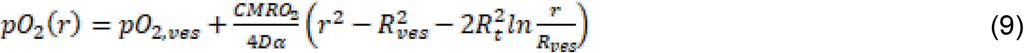

Similar to ODACITI model, this expression satisfies Eq. 4 for *R_ves_* ≤ r ≤ *R_t_*, However, the Krogh-Erlang model requires that O_2_ delivered by the vessel in the center is consumed by the tissue within the cylinder with the radius *R_t_*, which implies that O_2_ flux 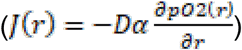 is equal to zero at r = *R_t_*. In the Krogh-Erlang model, the expression for the O_2_ flux (*J_KE_*(*r*)) for *R_ves_* ≤ r ≤ *R_t_*. is given by:

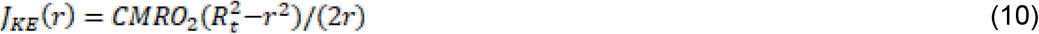

and total O_2_ flux through any cylindrical surface with length Δ*z* and radius r from the vessel is equal to the product of metabolic rate of O_2_ and volume of the tissue that remains outside the cylinder with radius r:

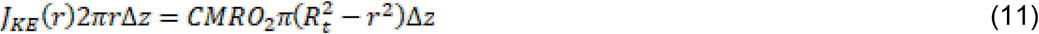

If we denote the O_2_ flux in ODACITI as *J_OD_*(*r*), total O_2_ flux through the wall of the central arteriole, for the arteriolar segment of length Δ*z*, is given by

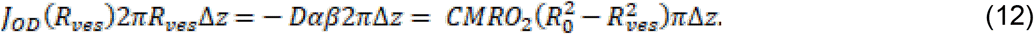

For any cylindrical surface with radius r between *R_ves_* and *R_t_*, the total O_2_ flux in ODACITI is given by

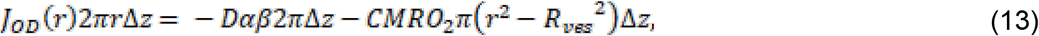

which represents a difference between the oxygen diffused from the vessel out 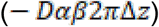 and the oxygen consumed in the inner tissue cylinder 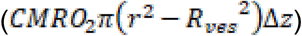. Finally, total O_2_ flux through a cylindrical surface with *r* > *R_t_* is constant:

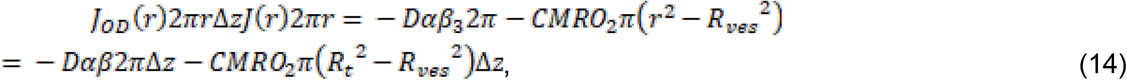

which is in agreement with the model assumption that for r>R_t_, CMRO_2_ is exactly equal to O_2_ delivery rate of capillary bed. In ODACITI, if oxygen flux at r=R_t_ is zero (*J_OD_*(*R_t_*) = 0), then *R*_0_ = *R_t_*, all oxygen delivered by the arteriole is consumed inside the r<R_t_ tissue volume (e.g., 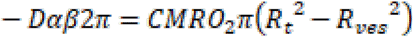) and ODACITI scales back exactly to the Krogh-Erlang model with additions that pO_2_ = const for r<R_ves_ and r>R_t_. However, in ODACITI it is possible that the radius where *J_OD_* (*r*) = 0 can be smaller than *R_t_*, which can better account for the influence of the capillary bed. In addition, this model allows to include more measured data points from small (r<R_ves_) and large radii (r>R_t_) into the fitting procedures as compared to using the Krogh-Erlang model.

### Segmentation of regions of interest

ODACITI describes pO_2_ gradients around diving arterioles for an ideal case of radial symmetry. In practice, however, capillary geometry around diving cortical arterioles is not perfectly radially symmetric. The model also assumes that pO_2_ decreases monotonically with increasing radial distance from the arteriole until reaching a certain stable level within the capillary bed. In reality, however, at certain distance pO_2_ may increase again due to presence of another highly oxygenated blood vessel that can be (a branch of) a diving arteriole or a post-arteriolar capillary. To mitigate this issue, we devised an algorithm for automated segmentation of a region of interest (ROI) for each acquired pO_2_ grid. First, tissue pO_2_ measurements were co-registered with the vascular anatomical images to localize the center of the arteriole. Next, 20 equally spaced radii were drawn in the territory around a center arteriole and a pO_2_ vector was interpolated as a function of the radial distance from the arteriole. These vectors were spatially filtered using the cubic smoothing spline *csaps* MATLAB function with smoothing parameter p=0.001 to reduce the effect of noise in the measurements. For each vector, we calculated the first spatial derivative 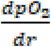 and included all pO_2_ data points with radii where 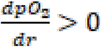. Finally, we added all pO_2_ measurements within the low 35% tail of each pO_2_ map to include capillary bed data points. Taken together, these steps resulted in exclusion of data points around additional oxygen sources within the capillary bed deviant from the model assumptions. Included pO_2_ data were then collapsed to generate a radial gradient for estimation of CMRO_2_ with the ODACITI model for each peri-arteriolar grid. When multiple acquisitions of the same plane were available, CMRO_2_ estimates were averaged.

### Synthetic data

To validate estimation of CMRO_2_ using ODACITI, we generated synthetic pO_2_ data by solving the Poisson equation for a given (space-invariant) value of CMRO_2_ and chosen geometry of vascular sources and measurements points. This was done numerically using the finite element software package FEniCS (Logg et al 2012) as described in our recent publication (Saetra et al 2020). Briefly, we solve the variational formulation of the Poisson equation where CMRO_2_ is constant, intravascular pO_2_ is fixed, and there is no pressure gradient at the vessel wall boundary. Then, if *V* is a space of test functions [*v*_1_, … *v_N_*] on the computational domain Ω, we can find pO_2_ such that 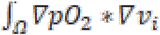 is equal to CMRO_2_ scaled by a number that depends on the test function *v_i_*. FEniCS provides the solution on an unstructured finite element mesh. Experimental data, in contrast, are measured on a Cartesian grid. Therefore, we transferred the synthetic data generated by FEniCS to a Cartesian grid similar to that used in our experiments. Additive Gaussian noise was implemented using the *normrnd* MATLAB function. For each synthetic data point in the grid, we drew a random number from a Gaussian distribution with the pO_2_ value at this point as the mean (μ) and standard deviation of σ=2. Afterwards, we replaced the pO_2_ value by (μ + σ).

### Statistics

Statistical analysis was performed using The R Project for Statistical Computing (www.r-project.org) using a linear mixed-effect model implemented in the lme4 package, where trends observed within a subject (e.g., the dependence of CMRO_2_ on depth) were treated as fixed effects, while the variability between subjects was considered as a random effect. Observations within an animal subject were considered dependent; observations between subjects, independent.

## Acknowledgements

We dedicate this paper to the memory of Andrei Vinogradov, a pioneer of mitochondrial energetics. We gratefully acknowledge support from the NIH (BRAIN Initiative R01MH111359, BRAIN Initiative U19NS123717, R01DA050159, R01NS108472, K99MH120053) and Swiss National Science Foundation (P2ZHP3_181568). Phosphorescent probe Oxyphor 2P was provided by the resource “Oxygen imaging by phosphorescence quenching”, supported by the grant U24 EB028941 from the National Institutes of Health, USA (directed by Dr. Sergei Vinogradov, University of Pennsylvania).

---

**Supplementary Figure 1.**
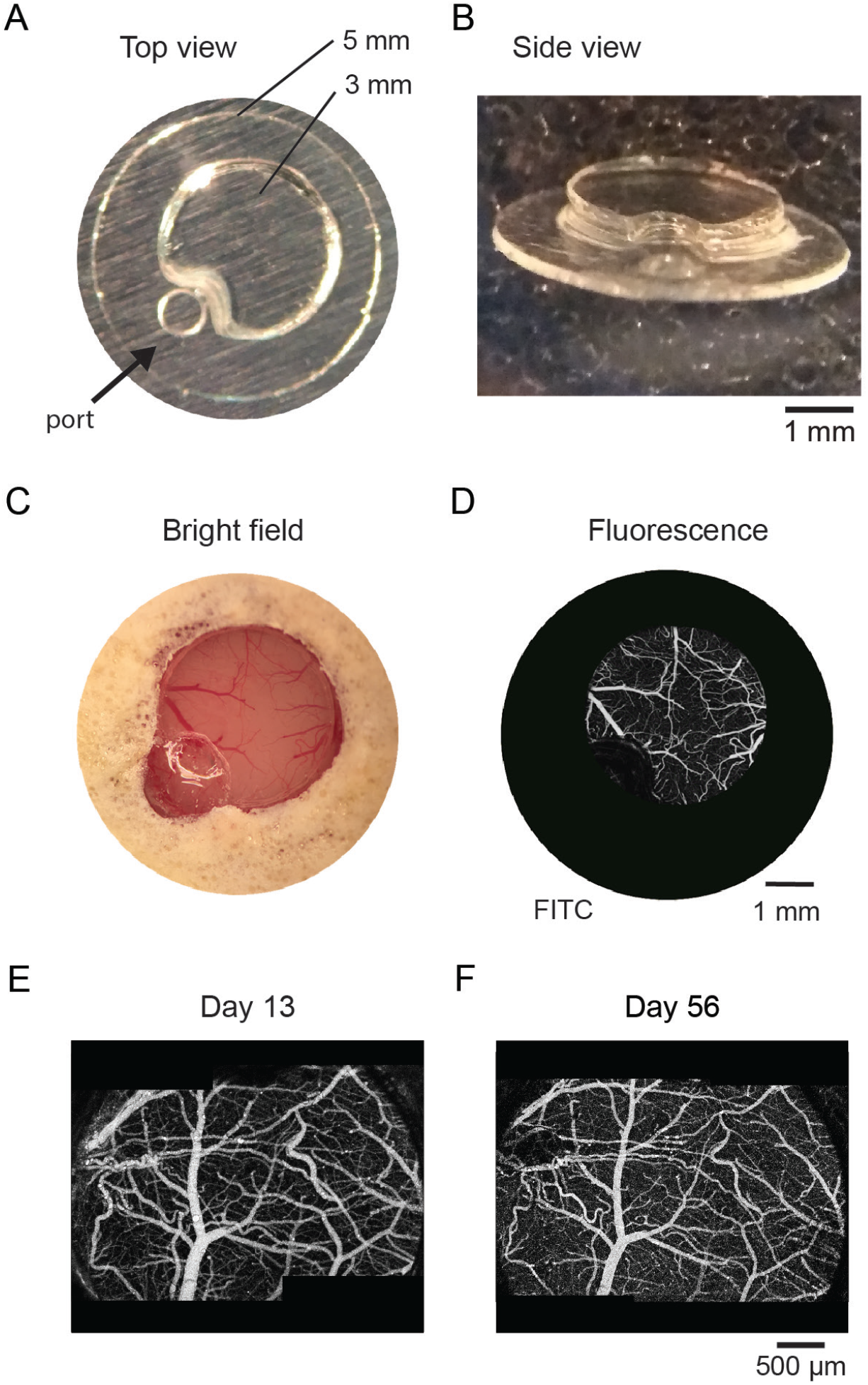
Optical window with a silicon access port. **A.** Top view of the window before filling the injection port with silicon. There are three 3-mm glass coverslips stacked together and glued to a 5-mm glass coverslip. The 3-mm stack is beveled to guide the pipette at an angle through the port. **B.** A side view of the window; the port is filled with silicon. **C.** A top view of an implanted window; surface blood vessels are visible under the glass. **D.** An image of surface vasculature within the same window calculated as a maximum intensity projection (MIP) of a 2-photon image stack 0-300 μm in depth using a 5x objective. Individual images were acquired every 10 μm. Fluorescence is due to intravascular FITC. **E.** Consecutive images obtained 43 days apart demonstrate stability of surface vasculature overtime.

**Supplementary Figure 2.**
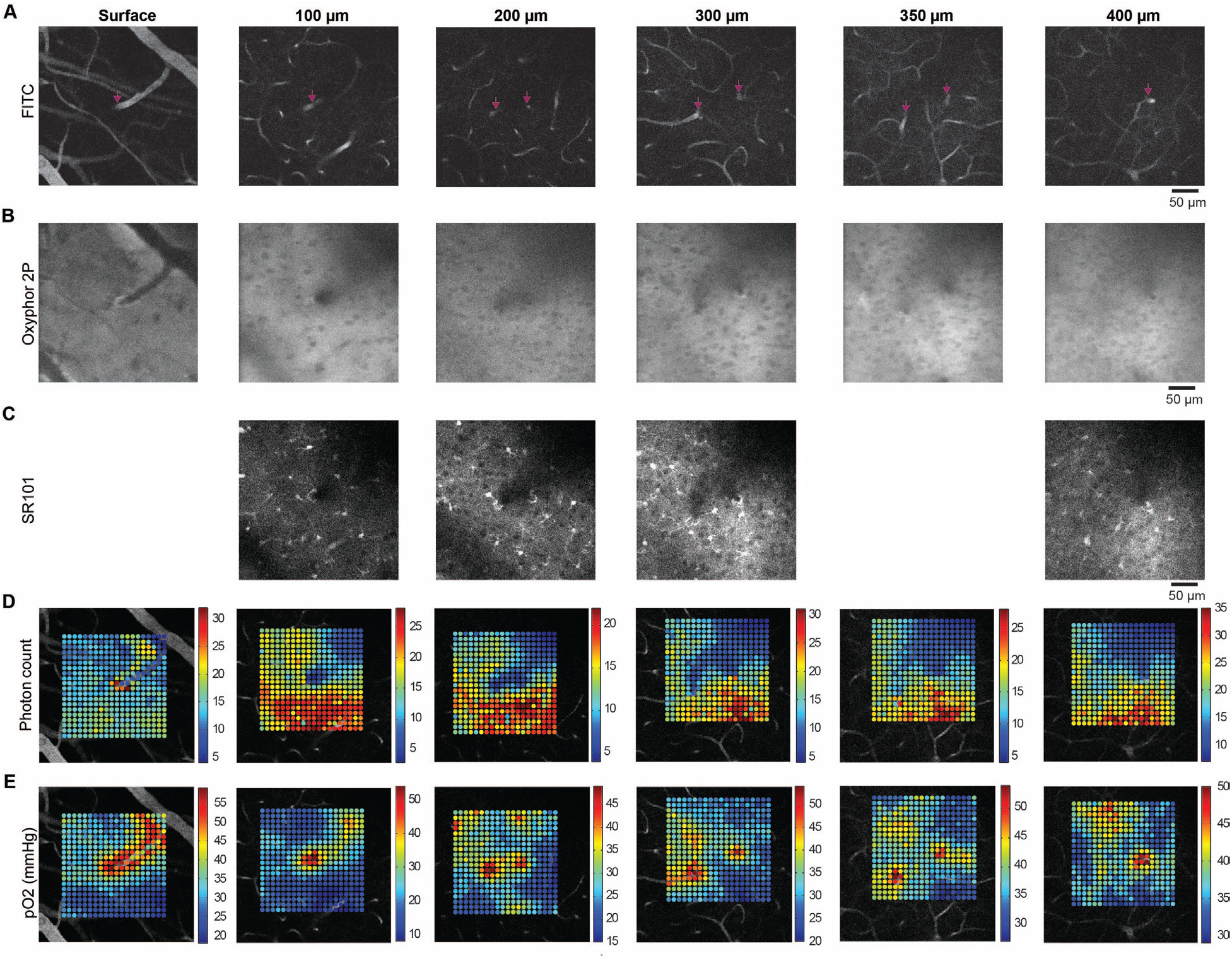
Example dataset for one arteriole across depths. **A.** Intravascular FITC images for six imaging planes (cortical surface, 100 μm, 200 μm, 300 μm, 350 μm, 400 μm). Arrows point to a diving arteriole that branches into two between 100 and 200 μm. **B.** Phosphorescence images for each of these planes. **C.** SR101 images; for two of the planes SR101 images were not acquired. **D.** A square measurement grid of 20×20 points for each of the imaging planes. Photon counts are superimposed on corresponding vascular FITC images. **E.** Calculated pO_2_ values superimposed on corresponding vascular FITC images.

**Supplementary Figure 3.**
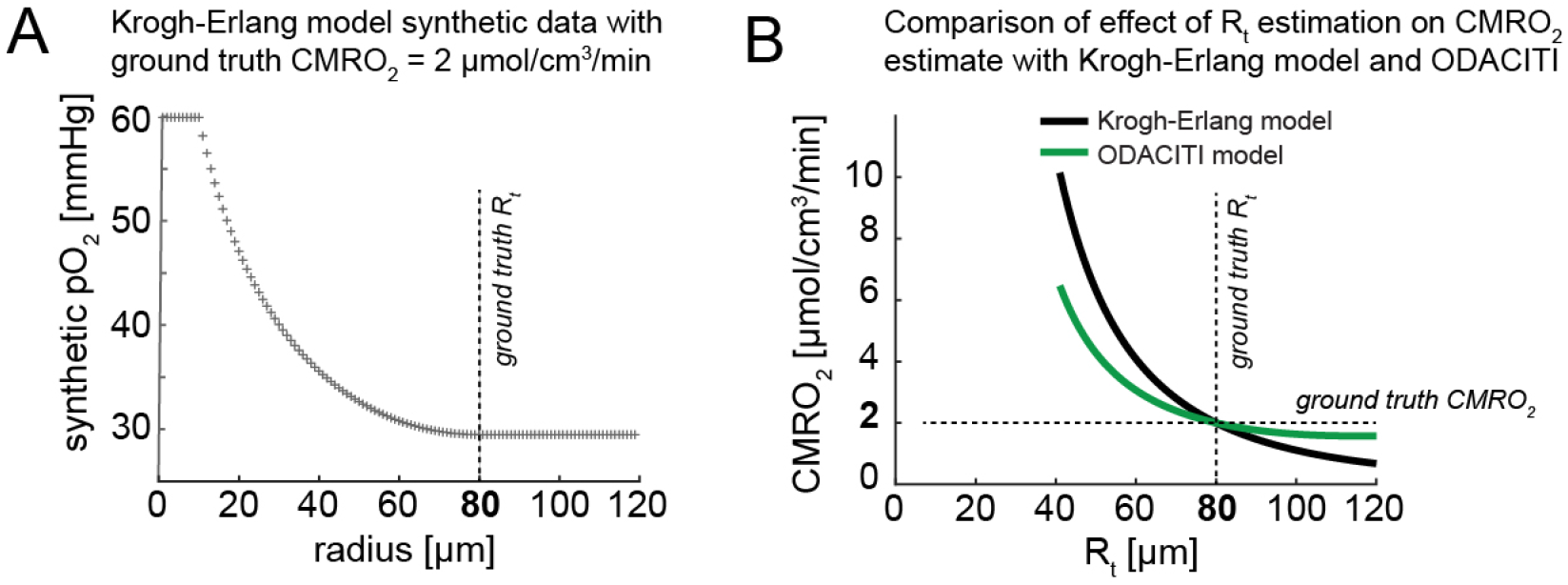
The effect of Rt on CMRO_2_ estimates. **A.** Synthetic pO_2_ data based on the Krogh-Erlang model (CMRO_2_ = 2 μmol cm-3 min-1, R_t_ = 80 μm, P_ves_ = 60 mmHg, R_ves_= 10 μm) are plotted against the distance from the arteriole. A constant pO_2_ is assumed for r> R_t_. **B.** The estimated CMRO_2_ after applying ODACITI (green) and the Krogh-Erlang model (black) on the synthetic data in panel A as a function of different assumed R_t_ values (the true R_t_ = 80 μm).

**Supplementary Figure 4.**
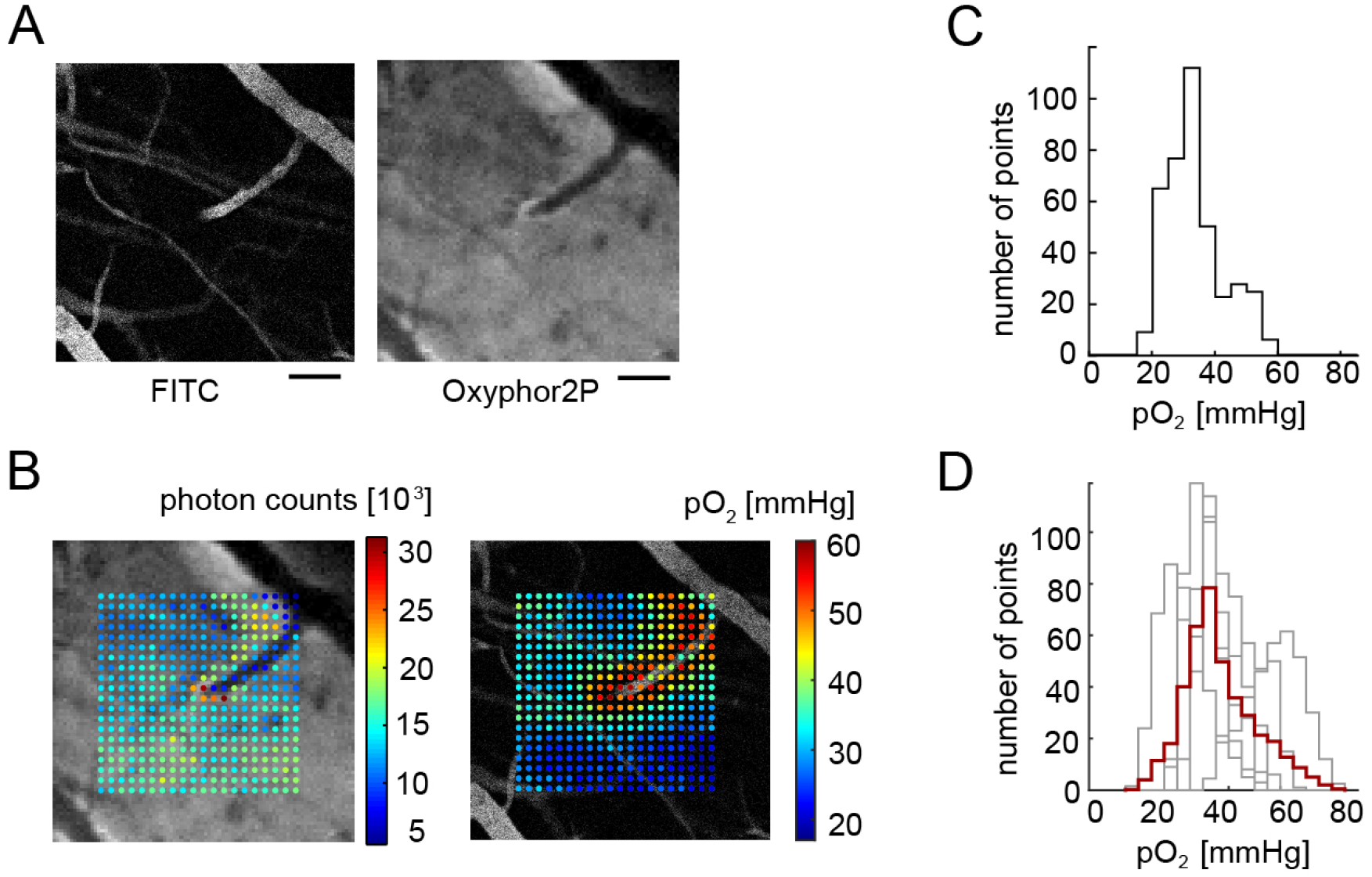
pO_2_ measurements at the cortical surface. **A.** Intravascular FITC image of a small surface arteriole (left) and a corresponding phosphorescence image (right). **B.** A square measurement grid overlaid on the FITC image from (A). Left: photon counts. Right: calculated pO_2_ values. Scale bar = 50 μm **C.** Histogram of pO_2_ values corresponding to (B). **D.** Overlaid histograms from each surface measurement plane (corresponding to individual arterioles, gray) and superimposed average (red).

**Supplementary Figure 5.**
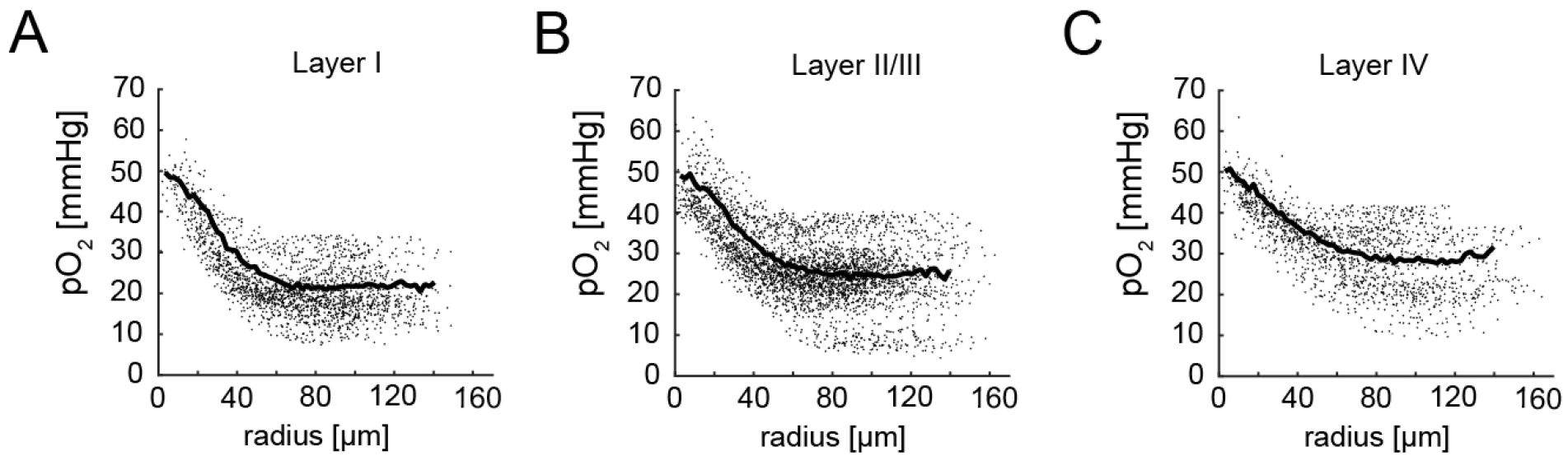
Cumulative radial pO_2_ profiles for each layer. **A.** All data points included in the CMRO_2_ estimation in Figure 3; the mean (calculated using 2.5 μm binning) is superimposed in thick black. **B.** The same as (A) for layer II/III. **C.** The same as (A) for layer IV.

**Supplementary Figure 6.**
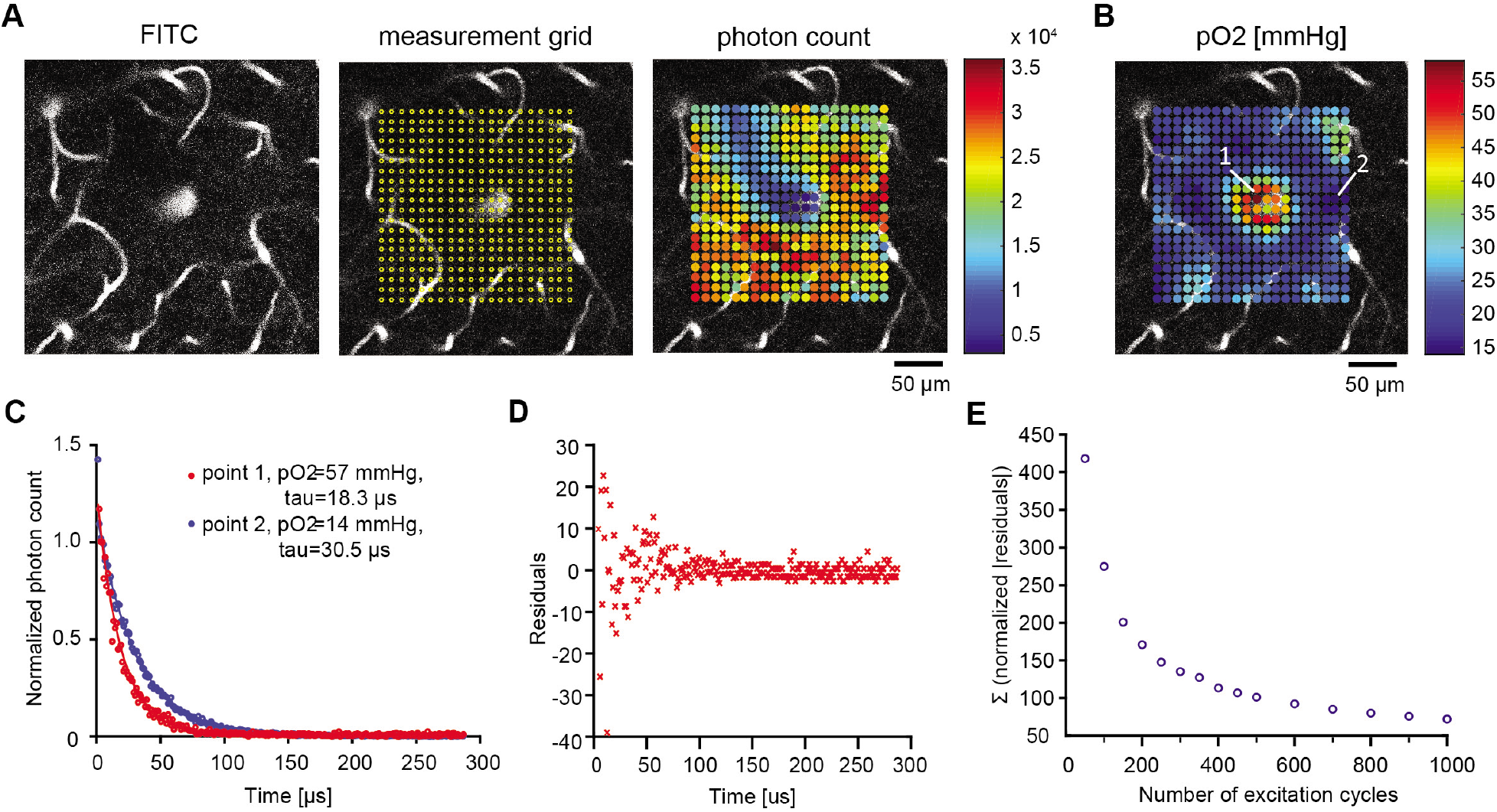
Accuracy of pO_2_ estimation as a function of number of excitation cycles. **A.** Left: an imaging plane 100 μm below the surface. Fluorescence is due to intravascular FITC. Middle: a grid of measurement points overlaid on the FITC image. Right: photon count overlaid on the FITC image. **B.** pO_2_ values overlaid on the FITC image from (A). **C.** Phosphorescence decays for two points labeled in (B). Data (points) and fit (lines) are overlaid. Lower pO_2_ corresponds to slower decay (blue). **D.** Residual error over time. **E.** The squared residual norm (output of *lsqnonlin)* plotted against the number of excitation cycles that were used to sum the photon counts.

